# Distinct but interacting functional filters of aridity and grazing shape Mediterranean mountain grasslands

**DOI:** 10.64898/2026.02.04.703801

**Authors:** Ioanna Nanopoulou, Fotiadis Georgios, Guillaume Delhaye, Konstantina Zografou, Vassiliki Kati, Charilaos Yiotis, Ioannis Tsiripidis, Anna Mastrogianni, Christina Kassara, Maria Petridou, Konstantina Nasiou, George C. Adamidis

## Abstract

Mediterranean mountain grasslands are ecosystems of high ecological and economic value. They are shaped by the dry and warm climate and land use, such as grazing, although the combined effects of both drivers remain poorly understood.

In this study, we analyzed shifts in functional composition in thirty-two plant communities in Mediterranean mountain grasslands of the Pindos Range (Greece) by measuring five plant functional traits related to resource acquisition in dominant plant species. We examined the adaptive value of each trait as well as community-level responses along a well-defined two-dimensional gradient of grazing intensity and aridity, using mixed models and functional diversity analyses, and tested whether individual species trait shifts are related to aridity and grazing intensity.

At the community level, aridity decreased plant height and leaf area whereas grazing only affected traits associated with tissue recovery such as high specific leaf area (SLA) and low community-weighted mean leaf dry matter content (LDMC). As aridity increased, plant height functional dispersion decreased. This convergence pattern indicates a shift towards more similar growth forms under arid conditions. Species-specific analysis indicated various responses of traits to the interaction of aridity and grazing that could not be detected using only community-level patterns.

Overall, our findings demonstrate that aridity and grazing act through separate functional axes at the community level, while their combined effects emerge through species-specific trait plasticity.

## INTRODUCTION

Mediterranean mountain grasslands are ecologically rich systems, sustaining high biodiversity and providing crucial ecosystem services, but changes in environmental conditions and land use result in a loss of biodiversity and ecosystem services (Ronchi & Brambilla 2025). These ecosystems are characterized by high seasonality and often have a long history of grazing (Grime 2001, Stanisci et al. 2020, Al Hajj et.al 2024). Mediterranean grasslands are among the most climate-sensitive ecosystems, and predicting their resistance to increasing aridity and land-use change requires an understanding of how climatic and land-use filters interact (Oñatibia et.al 2020, Zhu et.al 2024).

An effective approach to analyze potential interactions between abiotic and biotic factors is through a functional trait framework. Plant functional traits are individual-level measurable features that influence species fitness and provide information about species ecological strategies (Wright et al 2004). Trait-based approaches provide insights into how environmental gradients influence community composition and functional organization, linking species turnover and dominance patterns to underlying ecological mechanisms (Stanisci et al. 2020, Al Hajj et al. 2024). This approach is especially well-suited for assessing how several stressors act simultaneously because it identifies unique functional pathways through which abiotic and biotic filters influence community assembly.

Plant functional strategies of Mediterranean grasslands are influenced by aridity and grazing, but their effects may work through different ecological mechanisms. These systems have multiple strategies to cope with environmental stress, expressed as size and structure-related changes and coordinated shifts in leaf economic traits. A long-term stress, such as aridity, is more likely to affect plant architecture and morphology than tissue turnover (Valencia et. al 2015, Al Hajj et. al. 2024).

Increasing aridity limits water availability, so morphological and architectural traits that reduce water loss, such as lower plant sizes and compact growth forms, are favoured (Cingolani et al. 2005, Pakeman et al. 2009, Niu et al. 2016). Grazing, on the other hand, acts as a biotic filter through biomass removal and trampling, favouring plant species that can regenerate and recover quickly following defoliation. Such responses are linked with high specific leaf area (SLA) and low leaf dry matter content (LDMC), mainly under repeated or moderate grazing pressures (Díaz et al. 2007, Frennette-Dussault et al. 2012, McNaughton 1979, 1983). Because grazing is recurring rather than a constant resource limitation, it is likely to favor traits linked with quick tissue replacement and compensatory growth. Intense grazing may favor disturbance-tolerant species and constrain functional variation, while moderate grazing promotes compensatory growth and sustains functional diversity (Díaz et al. 2007, Rahmanian et al. 2019, Rakosy et al. 2022).

Different functional diversity metrics exist to provide insights into how environmental filters affect plant communities through interspecific trait variation (e.g. de Bello et al 2021). Community-weighted mean (CWM) trait values summarize dominant functional strategies within communities by weighting species traits based on their relative abundance, thereby capturing changes in functional composition generated by species turnover and dominance patterns (Laliberté & Legendre 2010, Muscarella & Uriarte 2016).

In addition, functional dispersion (FDis) quantifies the spread of functional strategies within communities and provides insights into community assembly processes by distinguishing between trait convergence driven by environmental filtering and trait divergence caused by niche complementarity (Laliberté & Legendre 2010; Valencia et al. 2015). CWM and FDis work complementarily to determine whether environmental gradients primarily shift average trait values, constrain functional diversity, or both, providing a strong framework for disentangling the effects of multiple stressors acting along different functional axes (de Bello et al. 2021).

Metrics at the community level, such as CWM and FDis, can reveal dominant strategies and functional structure but may not reveal individual species’ responses. Species in the same community can respond differently to the same environmental gradients, employing alternate functional strategies that cancel each other out when averaged at the community level (Albert et al. 2011, Violle et al. 2012, Siefert et al. 2015). Weak or missing community-level responses may indicate considerable variation in species-specific responses, rather than poor functional adjustment.

Species-specific trait responses help to interpret community-level patterns by showing how different species may adjust their strategies under different abiotic and biotic pressures. Interspecific variation in Mediterranean grasslands is expected to strongly influence community assembly and the distribution of functional strategies. This is more evident when environmental filters (such as aridity and grazing) act on different trait dimensions (Albert et al. 2011, Violle et al. 2012).

In this study, we use a trait-based approach to investigate the effects of grazing and aridity on the functional structure of Mediterranean mountain grasslands. We analyze community-weighted mean trait values, functional dispersion, and species-specific trait responses across a dual aridity and grazing intensity gradient to assess their influence on interspecific functional composition.

We focus on key traits related to plant size, resource acquisition, and tissue investment. We aim to investigate whether aridity and grazing act through shared or distinct functional strategies at the community level and to examine species-specific strategies as a response to the combined pressures of climate and land-use.

We specifically expect:

1. Community-level trait compositions and functional diversity to shift along aridity and grazing gradients. Specifically, aridity is expected to favor stress-tolerant strategies, while grazing is expected to favor species with fast recovery after biomass removal.
2. Strong interspecific differences in trait responses along the aridity and grazing gradients, with potential diverse species-specific interactions between stressors.

This integrated approach enables us to investigate how abiotic and biotic stressors may interact to shape the functioning of Mediterranean mountain grasslands under increasing climate and land-use change.

## MATERIALS AND METHODS

### Sites description and plant communities

Sixty-four 1m x 1m plots were sampled in June 2024, across thirty-two sites located within the Pindos mountain range, 21 of which were situated inside Natura 2000 protected areas (Fig. 1). All study sites were above the treeline (1,470–1,850 m a.s.l.), characterized by gentle slopes (<30°), and were chosen to maximize the variability of both the aridity and grazing intensity in the grasslands, creating a system of two interacting stress gradients. At each site, one transect was established, and two replicate plots were placed along it, separated by 200 m. Species abundance was recorded through visual estimation of percentage (%) cover for each species, following the Braun-Blanquet scale (Sutherland 1997). Species were identified in situ and, when necessary, in the lab using a dissecting microscope stereoscope.

**Figure 1.**
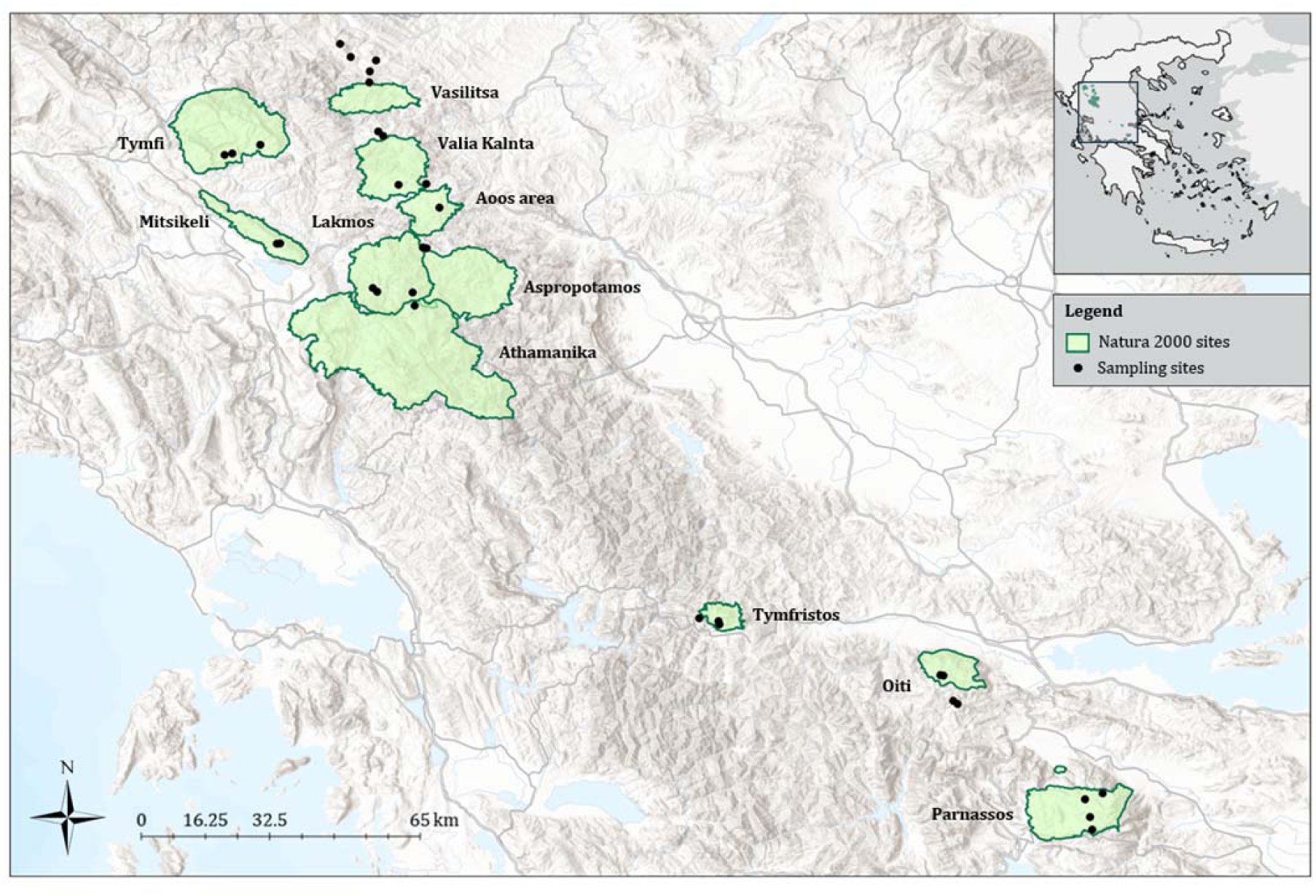
Sampling sites (32 sites) across the Pindos mountain range, indicating sites falling within the Natura 2000 network of protected areas (green polygons).

### Aridity and grazing variables

To capture variation in aridity among sites, we calculated the normalized difference infrared index (NDII hereafter) and the normalized difference vegetation index (NDVI hereafter) for the period of May - August of 2024. These indices are sensitive to water and chlorophyll content as well as to the vegetation structure of the study sites (Tucker 1979, Jiang et al. 2006, Yilmaz et al. 2008). Grazing intensity was estimated based on semi-structured interviews with livestock farmers using the pastures overlapping our study sites. For each herd, we recorded herd size, livestock type, and age classes, as well as the grazing period. Grazing duration was converted to grazing months, calculated as the total number of grazing days divided by 30.44 (average number of days per month). Livestock numbers were converted into Livestock Units (LSU) using standard Eurostat coefficients (Eurostat 2020; i.e. cattle >2 years = 0.8, cattle 1–2 years = 0.7, cattle <1 year = 0.4, sheep = 0.1, goat = 0.1, equidae = 0.8). For each study site, livestock units were summed across all herds grazing the site to derive total grazing pressure.

Based on these data, different LSU-based indices were evaluated to examine their correlation structure and suitability as a single grazing pressure metric. Based on these evaluations, total livestock units expressed in grazing months (LSU.MN.Total, hereafter), incorporating all herd types and the full grazing period, were selected to represent grazing intensity in our analysis, as they best capture the dominant, cumulative pressure in the study area.

### Trait measurements

Five functional traits related to plant growth, nutrient use, stress tolerance and competitive ability were measured on fully expanded leaves of the most abundant species across our study sites, for a minimum of two dominant species per plot representing on average 81% of the total cover (60%–97%). Traits were measured on a minimum of three individuals per species at the site level, following standardized procedures (Pérez-Harguindeguy et al., 2013). Leaves were placed flat on a white background, photographed, and leaf area was estimated using ImageJ software.

Specific leaf area (SLA) was calculated as the ratio of the water-saturated leaf area to the leaf dry mass and leaf dry matter content (LDMC) was determined as the ratio of leaf dry mass to water-saturated fresh mass. Leaf thickness (LT) was estimated to the nearest 0.01mm using a caliper. Plant height (H) was measured as the maximum vertical distance (cm) from the ground level to the highest point of the main photosynthetic tissue of a mature individual keeping the plant in its natural position. For abundant species present in the plots whose traits could not be measured at the time of sampling, trait values were obtained from the TRY database in order to increase coverage of dominant species.

### Statistical analysis

To understand the relationships between predictor variables, we performed a principal component analysis (PCA), with NDVI, NDII and LSU.MN.Total. Based on the variable loadings, the first axis (PC1) was interpreted as an aridity gradient, and the second axis (PC2) was interpreted as an independent grazing intensity gradient (Figure 2).

**Figure 2:**
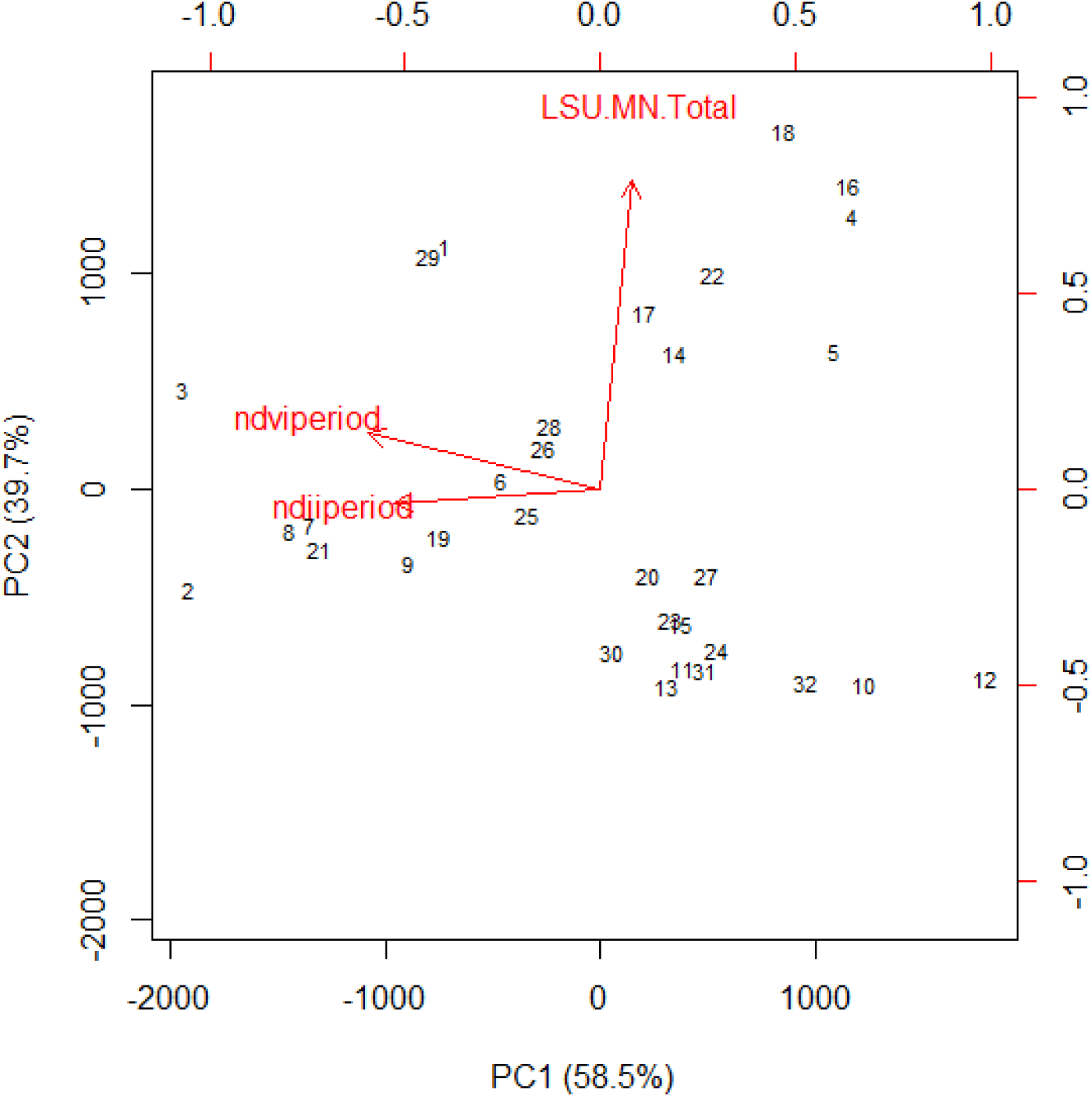
Principal components analysis combining data on aridity (ndviprtiod, ndiiperiod) and grazing (LSU.MN.Total) intensity from our 32 sites across Pindos Mountain range. The first two principal components explained 58.5% and 39.7% of the total variance, respectively, with PC1 accounting for the largest proportion of variation.

### Community trait composition

Species composition data were pooled per site prior to analyses. While the conventional approach of regressing community-weighted mean trait values along environmental gradients (CWMr) is informative for evaluating community-level effects on ecosystem processes, the adaptive value of traits should be assessed using multilevel models including species as random effects (Lepš and de Bello 2023, Jamil et al., 2013, ter Braak, 2019). In this study, we used both approaches, as they provide complementary information regarding the future distribution of species and the ecosystem processes.

To evaluate trait adaptability, we implemented the MLM3 approach described by ter Braak (2019). This method was selected for its strong statistical properties, particularly its ability to minimize Type I error while maintaining high sensitivity compared to alternative multilevel designs. This method uses a multilevel linear model to estimate the abundance of each species as a function of environmental gradients (here, aridity and grazing intensity), trait values, and their interactions, while accounting for differences between species and sites through random effects. To prevent low abundance values from being converted into artificial zeros, abundance data were rounded to the nearest upper integer using a ceiling function. The final mixed-effects model included the interaction between mean species trait values and the combined aridity-grazing gradient as a fixed effect. Two random components were incorporated: one capturing trait variation among sites and another representing species-specific responses to the aridity-grazing gradient. The response variable was modeled using a Poisson distribution.

To quantify dominant trait patterns and their potential effects on ecosystem functioning, we calculated the Community Weighted Mean (CWM) for each trait within each sampling site (Lavorel & Garnier, 2002). We also explored the relative contribution of various community assembly mechanisms using the functional dispersion index (FDis) (Laliberté & Legendre, 2010), which quantifies the mean distance of each species from the centroid of a multi-trait space, weighted by their relative abundance. Because multivariate functional diversity metrics may mask patterns associated to certain axes of functional variation (Butterfield & Suding, 2013), we additionally calculated FDis for each trait separately. As functional diversity measures are sensitive to the total number of species present, we performed a null model approach to compute the standardized effect size of FDis (sesFDis), following Delhaye et al. (2020; 2024). For each site, we produced a null distribution of FDis by randomly assigning species abundances (n=500 times) and calculated the mean and standard deviation of this distribution. The sesFDis was calculated using the formula: 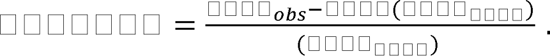

Both CWM and FDis were calculated using species-level median trait values. Relationships between FDis and CWM with the stress gradients were analyzed using multilevel linear models, with mountains included as a random effect.

### Species-specific trait analysis

Species-specific responses to the interaction of aridity and grazing intensity were analyzed using linear mixed-effects models. Each functional trait was fitted separately. The analysis was limited to species recorded in the field that had sufficient trait measurements. We only used species with at least three observations per trait for all studied traits.

Before model fitting, traits were standardized, and aridity and grazing were scaled. We used mixed-effects models with fixed effects for the interaction of aridity and grazing. Species were included as random effects, resulting in random intercepts and slopes for aridity, grazing, and their interaction.

Species-specific aridity by grazing interactions were evaluated based on the random effect estimates and their associated confidence intervals. If the confidence intervals of the interaction term didn’t overlap zero, species considered to have interacting responses. We calculated the proportion of species showing significant interactions.

We initially fitted generalized additive models (GAMs) for the mixed-effects analyses. However, the estimated degrees of freedom (edf) indicated no evidence of nonlinear relationships, and diagnostic plots likewise suggested linear associations among variables. Consequently, we proceeded with linear mixed-effects models. Spatial autocorrelation was assessed using DHARMa residual diagnostics (Hartig.F. 2024). When spatial autocorrelation was detected, it was addressed either by refitting the models within a GAM framework without smooth terms, thereby preserving linearity, or by applying generalized least squares (GLS) models. Data processing and visualization were carried out in R (R Core Team, 2025) using RStudio (RStudio Team, 2025) and the tidyverse package suite (Wickham et al., 2019).

## RESULTS

### Changes in trait composition towards stress tolerance along stress gradients

Along a gradient of increasing aridity, species characterized by higher leaf thickness (β = 0.25, p = 0.00), lower leaf area (β = −0.24, p = 0.01), and shorter stature (β = −0.24, p = 0.00) become more abundant (Figure 3A).

**Figure 3.**
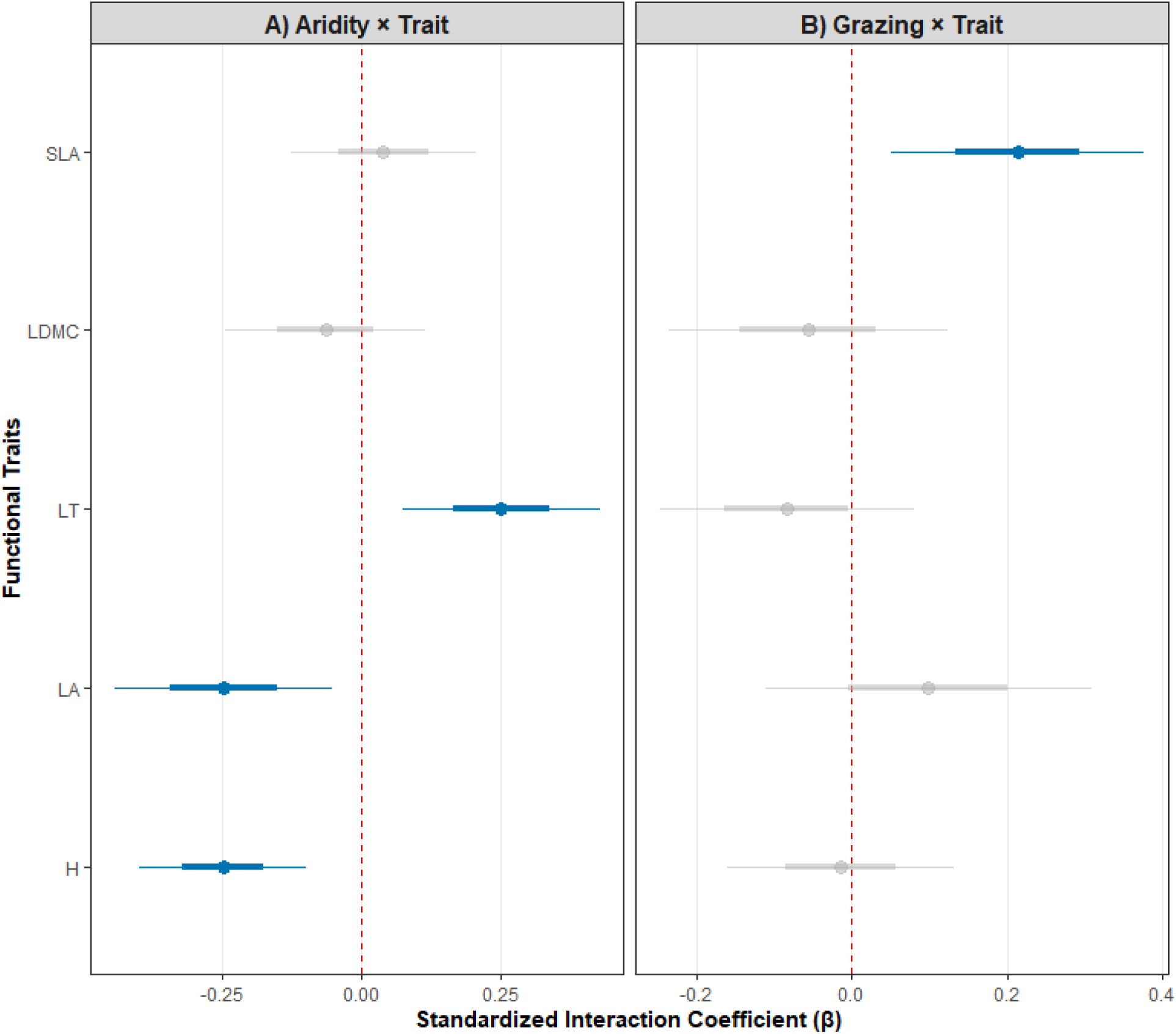
Forest plots on the effects of trait–environment relationships across the whole community, showing estimated coefficients (β) for the interaction between aridity-grazing and species trait values derived from generalized linear mixed models. Points represent coefficient estimates, thick horizontal bars indicate 66% confidence intervals, and thin horizontal bars indicate 95% confidence intervals. The blue line shows statistically supported effects (p = 0.05), whereas the grey line indicates unsupported effects.

The aridity gradient didn’t show consistent trait-environment relationships for all traits. There was not a significant response for SLA and LDMC for aridity (Figure 3A). Changes in species abundance with increasing aridity, at the community level, are mostly linked to plant size and leaf structural traits rather than leaf economic traits.

Grazing showed a more restricted effect. Only SLA was associated with grazing, with species with high SLA values increasing their abundance with increasing grazing intensity (β = 0.21, p = 0.01) (Figure 3B), pointing to a shift towards more acquisitive, fast-growing strategies in response to increasing grazing pressure.

Overall, aridity had an impact on several traits such as plant height, leaf area and leaf thickness, whereas grazing effect was only restricted to SLA. Notably, no trait responded to both aridity and grazing interaction, indicating a distinction of abiotic and biotic filtering mechanisms.

Community-weighted means (CWM) showed patterns similar to the previous models for most traits. Increasing aridity significantly reduced leaf area (slope=-0.48, p=0.00) and height of the communities (slope=-0.40, p=0.01; Figure 4A). In contrast, the communities exhibited a clear decrease in LDMC with increasing grazing intensity (slope=-0.40, p=0.05; Figure 4B).

**Figure 4.**
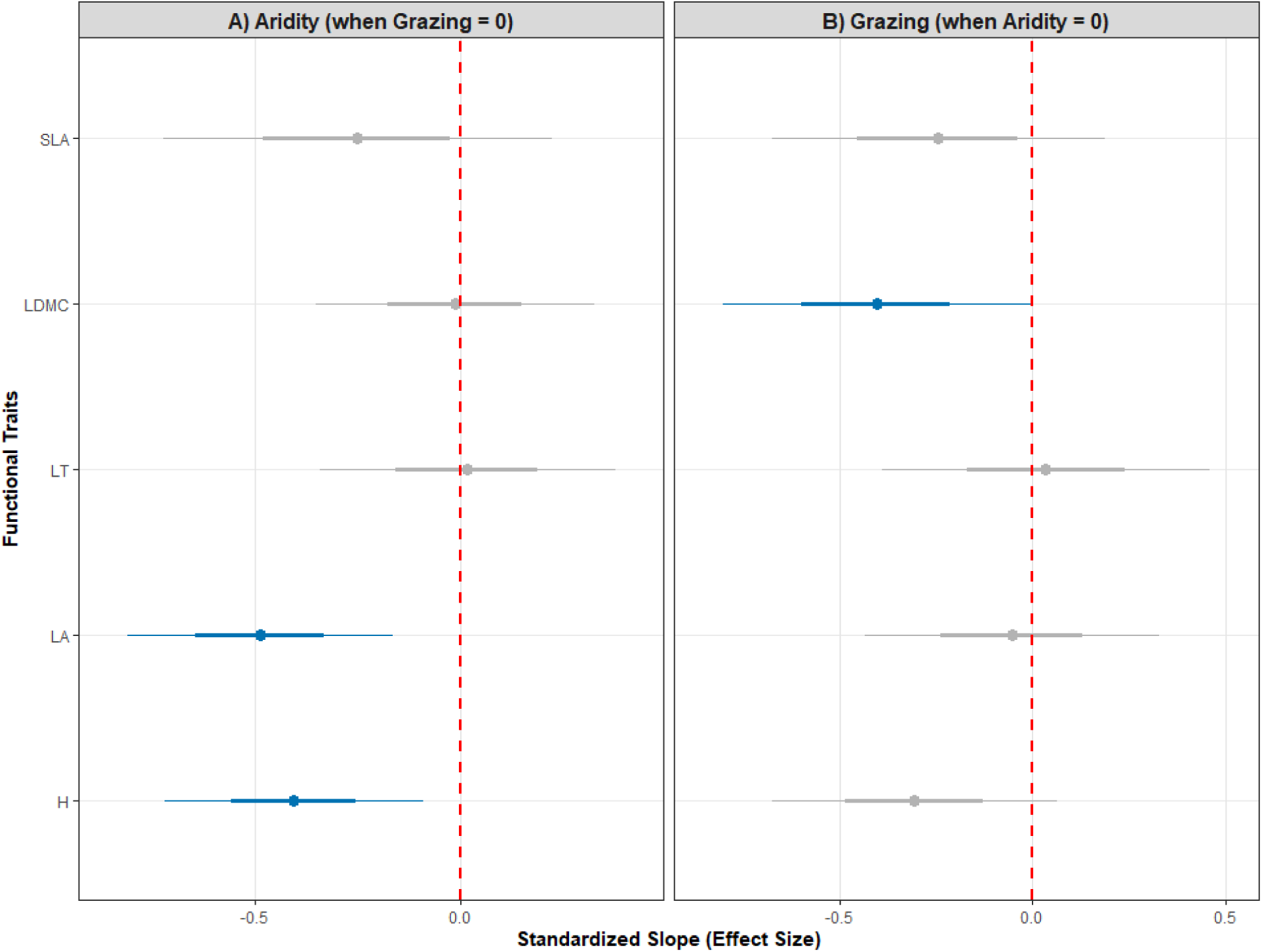
Forest plot of standardized slopes from linear models (LMMs/GAMs). The dots represent the mean estimates, while the thick and thin lines represent the 66% and 95% confidence intervals, respectively. Blue indicates interspecific diversity. Intervals that do not intersect the dotted line (zero) indicate statistically significant effects (p < 0.05).The grey line indicates non-supported effects.

Species abundance and community-weighted mean results indicate that grazing intensity leads to species with higher SLA values and smaller LDMC. When taken together, this shows that grazing promotes functional strategies with quick tissue recovery and high renewal capacity.

Regarding the functional dispersion, the multivariate metric did not show any significant shift along the aridity-grazing gradient. Univariate functional dispersion showed a clear response only for plant height. Specifically, functional dispersion of plant height (FDis) decreased strongly as aridity increased (slope=-0.47, p=0.00; Figure 5). This trend indicates that as aridity increases, the range of height values within the community decreases, indicating a higher homogeneity in this trait. In contrast, no significant relationships were found between grazing intensity and height functional dispersion, indicating that grazing’s effect on height variation differs between communities.

**Figure 5.**
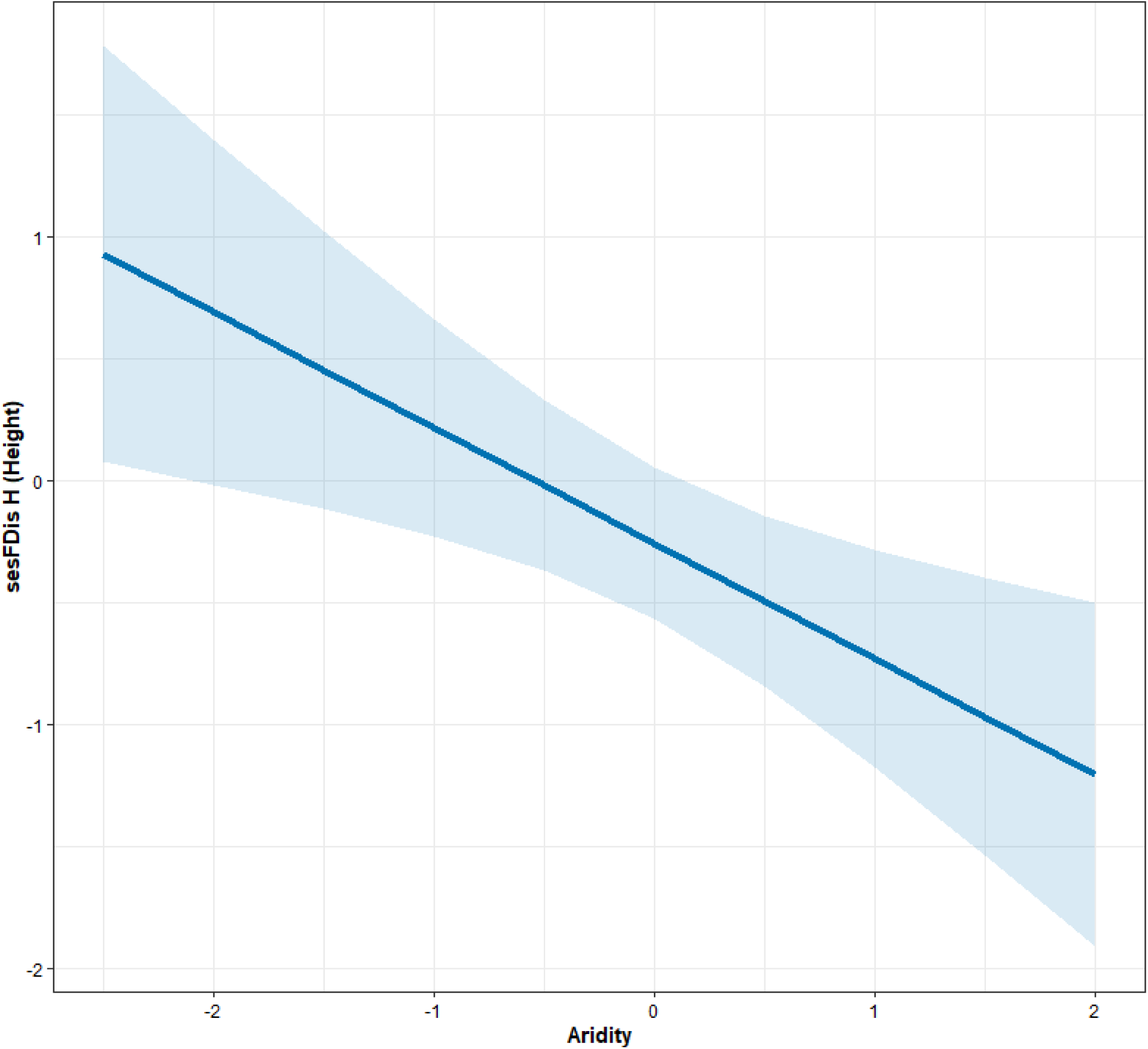
Predicted response of functional dispersion (FDis) for plant height along the aridity gradient. The line represents values for the interspecific component (blue), with the shaded area indicating a 95% confidence interval.

### Species-Specific Responses

Our species-specific analysis of trait shifts along the studied gradients revealed strong and diverse responses, even when overall fixed effects were weak. The interaction between aridity and grazing intensity was statistically significant for plant height in 44.7% of the studied species, whereas substantially fewer species showed significant interactions for leaf traits, including leaf area (2.6%), LDMC (18.4%), leaf thickness (7.9%), and SLA (10.5%), indicating that the combined influence of aridity and grazing was expressed primarily through plant height, while leaf-level traits were less frequently affected by their interaction. Among species exhibiting significant interactions with the aridity-grazing gradients, plant height decreased in 52.9% (9 species) and increased in 47.1% (8 species). Regarding the leaf functional traits, leaf area increased in the only species exhibiting a significant interaction. LDMC decreased in one species and increased in two species, whereas SLA responses were evenly split, with two species showing decreases and 2 species showing increases (Table S1).

For clarity, Figure 6 presents only species exhibiting significant interactions for more than one trait, whereas full model outputs for all species are presented in Table S1.

**Figure 6.**
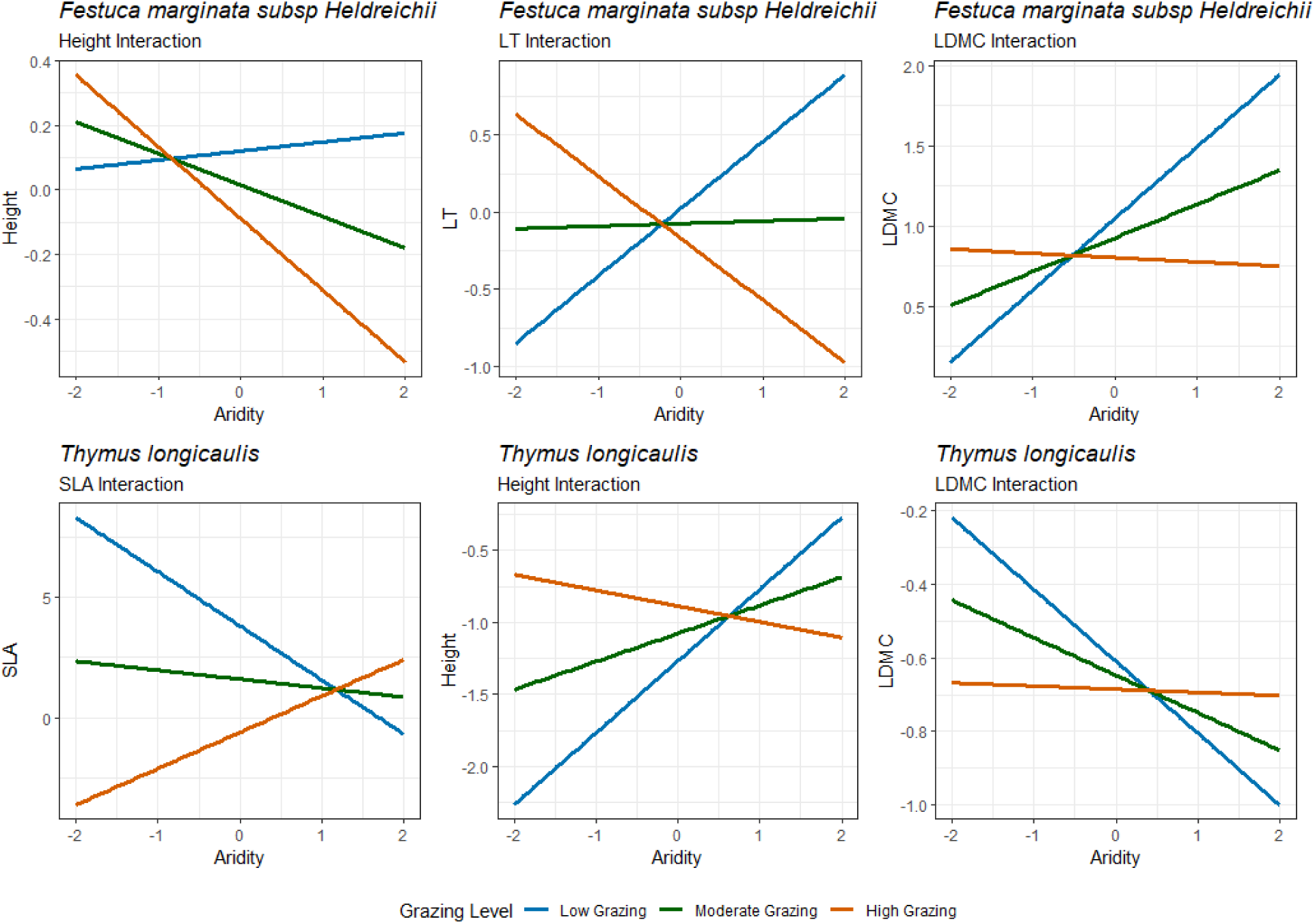
Species-specific trait responses to the aridity gradient under three levels of grazing intensity. Panels illustrate the significant interaction effects (random slopes) between aridity and grazing for selected species (Festuca marginata subsp. heldreichii, and Thymus longicaulis). Lines represent model predictions for low (blue), moderate (green), and high (orange) grazing levels.

*Festuca marginata subsp. heldreichii* showed significant interactions among various traits. In low grazing pressure as aridity increased, plant height also increased, whereas at moderate and high grazing, the species tended to decrease its stature with increasing aridity. Leaf thickness increased with increasing aridity at low grazing pressure, but at high grazing, it decreased. At moderate grazing, leaf thickness was not affected by aridity. Also, LDMC increased at low and moderate grazing and decreased at high grazing with increasing aridity.

Several traits seemed to have significant interactions for *Thymus longicaulis* too. SLA decreased with increasing aridity under low grazing, slightly decreased at moderate grazing, but increased under high grazing. With increasing aridity at low and moderate grazing, taller plants were promoted, but at high grazing pressure, plant height decreased. LDMC decreased with aridity under low and moderate grazing pressures, whereas in moderate grazing, it appeared to be stable.

Overall, these responses demonstrated solid and accurate correlations between aridity and grazing across various functional traits. Significantly, *T. longicaulis* and *F. marginata subs. heldreichi* both showed coordinated changes in height and functional leaf traits, indicating a reorganization of growth and leaf economic strategies under the interaction of aridity and grazing intensity.

## DISCUSSION

The impacts of biotic and abiotic filters on Mediterranean grasslands are thoroughly documented (de Bello et al. 2005, Bernard-Verdier et al. 2012), but their combined effect on the functional structure of mountain communities is less known. This study investigated how aridity and grazing intensity affect community-weighted mean trait values, functional dispersion, and species-specific trait responses.

Our results indicate that aridity and grazing act through distinct trait axes at the community level, while interacting through species specific plastic responses. Aridity primarily limited plant size and morphological traits such as leaf area and leaf thickness, whereas grazing effects were more specifically reflected through traits linked with tissue turnover and recovery, such as SLA and LDMC. Despite weak fixed effects at the community level, substantial species-specific responses demonstrated strong context-dependent plasticity, underscoring the significance of intraspecific variation in regulating plant responses to associated climate and land-use stressors (Albert et al. 2011). These findings provide fresh insights into the processes that maintain functional variability in Mediterranean mountain grasslands despite rising environmental stress (Carmona et al. 2012, Al Hajj et al. 2024, Zhu et al. 2024).

### Functional responses along the aridity–grazing gradient

Our analyses on the functional composition of the communities reveal that aridity and grazing work independently in Mediterranean mountain grasslands, influencing a number of plant survival strategies. Aridity mainly reduced the abundance of tall species with increased leaf area while increasing the abundance of species with high leaf thickness. This indicates that in more arid environments, plants adopt smaller and more compact forms of growth (Cingolani et al. 2005; Pakeman et al. 2009; Niu et al. 2016). Aridity, on the other hand, had no significant effect on SLA and LDMC at the community level. This separate effect on size-related traits and leaf economic traits suggests that aridity filtering in Mediterranean mountain grasslands may be driven primarily by morphological and architectural constraints, rather than through coordinated shifts in leaf economic traits (Cingolani et al. 2005, Pakeman et al. 2009).

Conversely, the effects of grazing are consistent with trait syndromes of acquisitive functional strategies. Increased grazing resulted in increased abundance of species with higher values of SLA and lower values of LDMC in the communities, indicating a shift towards fast-growing species with quick turnover strategies and low investment in dense leaf tissues. These patterns line up with compensatory growth responses, in which plants prioritize rapid tissue replacement above structural resistance after ongoing defoliation (McNaughton 1983, Cingolani et al. 2005, Zhao et al. 2008).

Grazing was trait-specific, with no consistent grazing-related responses observed for plant height, leaf area, or leaf thickness. Additionally, no trait was linked to both aridity and grazing interaction. This indicates that abiotic and biotic filters work through largely distinct pathways, shaping functional composition along multiple trait axes rather than an integrated stress-response syndrome (de Bello et al. 2005, Bernard-Verdier et al. 2012).

### Functional dispersion (FDis) and community assembly processes

Aridity and grazing shape the functional structure of Mediterranean mountain grasslands not only by affecting the mean trait values but also by altering the diversity of functional strategies (FDis). Functional dispersion captures the balance between environmental filtering and niche complementarity, providing insights into community assembly mechanisms (Laliberté & Legendre 2010, Weiher et al. 2011). In our study, functional dispersion decreased considerably with increasing aridity only for plant height, revealing a restriction in the range of height values represented within communities along the aridity gradient. This indicates a convergence of plant statures under arid environments. This broad convergence implies that aridity imposes a severe and consistent restriction on plant size, favoring growth forms with low water loss and maintenance costs (Cingolani et al. 2005, Pakeman et al. 2009, Niu et al. 2016). Grazing had no significant effect on functional dispersion of plant height, neither did aridity or grazing impact functional dispersion of leaf-related traits.

### Different roles of intraspecific variation and species replacement under abiotic and biotic filtering

Mixed-effects models revealed substantial species-specific responses to both stress gradients. The absence of strong fixed effects may reflect contrasting responses among dominant species, whose opposing strategies may cancel each other out at the community level, thereby masking plastic responses when averaged across species.

*F. marginata subsp. heldreichii* exhibited significant plasticity in plant height, leaf thickness, and LDMC, indicating context-dependent adaptation to grazing and aridity pressures (Henn et al. 2018). Plant height varies with grazing intensity, indicating a response to herbivory. Lower stature under high grazing pressure reduces exposure to grazers and mechanical damage while facilitating recovery through protected basal meristems (McNaughton 1983; Coughenour 1985, Milchunas et al. 1988, Díaz et al. 2007).

Leaf-level trait responses show a balance of structural investment and recovery capacity. Investment in dense, robust tissues that improve resilience to drought through better water retention and longer leaf lifespan is consistent with higher LDMC and thicker leaves under low grazing intensity (Wright et al. 2004, Pérez-Harguindeguy et al. 2013, Pakeman et al. 2009). As grazing pressure increases, however, structural investments become more limited, and reductions in LDMC lead to a shift towards lower-cost tissues and faster turnover strategies that promote rapid regrowth after defoliation (Díaz et al. 2007, Pérez-Harguindeguy et al. 2013, Al Hajj et al. 2024).

*T. longicaulis*, on the contrary, showed significant changes in functional strategy that were highly influenced by grazing intensity, showing context-dependent plasticity (Albert et al. 2011, Violle et al. 2012, Siefert et al. 2015). Low grazing pressure promoted plants with reduced SLA and LDMC values and increased plant height, along the aridity gradient, indicating a strategy that prioritizes tissue longevity over tissue recovery (Wright et al. 2004, Pérez-Harguindeguy et al. 2013, Reich 2014). Such trait combinations of SLA and height are frequently connected with climatic stress tolerance, where reducing resource loss and preserving durable tissues can be favorable in dry environments (Pakeman et al. 2009).

In alignment with our results, Zheng et al. (2024) showed that arid areas without grazing pressure (control areas) supported species with lower LDMC values in relation to highly grazed areas. They showed that in harsh environments, intense grazing acts as an extra pressure, which forces plants to invest in more dense and resilient tissues (S strategy), making ungrazed areas less conservative. Also, Frenette-Dussault et al. (2012) note that rising aridity may act as a filter that selects species with low LDMC as a stress-avoidant strategy to deal with water limitation independently of grazing pressures.

Under heavy grazing pressure, however, this strategy was reversed. As aridity increased, individuals became shorter with higher SLA values, indicating a change toward more acquisitive strategies that prioritize rapid metabolism, fast tissue turnover, and improved regrowth ability (Wright et al. 2004, Díaz et al. 2007, Pérez-Harguindeguy et al., 2013). This change is consistent with compensatory responses to repeated defoliation, in which rapid tissue replacement takes preference over investment in structurally costly tissues (McNaughton 1983, Coughenour 1985, Zhao et al. 2008).

Grazing-mediated modulation of drought-related trait expression is consistent with evidence that trait-environment interactions can vary under coupled pressures, resulting in diverse strategies across grazing contexts (Fajardo & Siefert 2018, Oñatibia et al. 2020, Zhang et al. 2023).

The opposing and complementing responses of *F. marginata subsp. heldreichii* and *T. longicaulis* show how species-specific plasticity contributes to functional variability in Mediterranean mountain grasslands. *F. marginata subsp. heldreichii* adapted through coordinated changes in plant architecture and tissue-level investment to balance structural resilience under low grazing with disturbance tolerance as grazing intensity increased. On the other hand, when aridity increased, *T. longicaulis* showed clear changes in resource-use strategy, moving from conservative trait syndromes under low grazing to more acquisitive, quick-turnover strategies under high grazing pressure.

## CONCLUSIONS

Our results reveal that aridity and grazing influence the functional structure of Mediterranean mountain grasslands by independent but interrelated processes. While their effects are mostly reflected along separate trait axes at the community level, their combined influence emerges by species-specific adjustment mechanisms.

These findings show that grassland sustainability under concurrent climatic and land-use stresses may rely on the coexistence of parallel functional strategies, where compensatory responses to grazing coexist with structural and architectural shifts to aridity. Future research combining trait-based techniques with long-term monitoring and experimental manipulation is needed to determine how such complementary strategies affect ecosystem functioning under increasing environmental stress.

## Author contributions

The manuscript was originally drafted by IN and GCA. Data analysis conducted by IN, GCA and GD. Field sampling was performed by IN, GCA, GF, CY, KZ, VK and KN. CK and MP collected data and calculated the aridity and grazing intensity indices. IT and AM shared functional trait data. All co-authors contributed to the interpretation of results, manuscript revisions, and approved the final version for submission.

