## Supplementary Material for "Distinct but interacting functional filters of aridity and grazing shape Mediterranean mountain grasslands"

| Species | SLA | Height | LA | LT | LDMC |
| --- | --- | --- | --- | --- | --- |
| *Achillea fraasii* | -0.16 (-0.56 : 0.24) | -0.05 (-0.1 : 0.01) | 0.07 (-0.17 : 0.32) | 0.12 (-0.14 : 0.38) | -0.01 (-0.09 : 0.06) |
| *Agrostis castellana* | -0.16 (-0.67 : 0.35) | 0.03 (-0.09 : 0.16) | -0.06 (-0.31 : 0.2) | -0.08 (-0.66 : 0.5) | -0.08 (-0.19 : 0.03) |
| *Anthemis arvensis* | 0.76 (-0.28 : 1.81) | -0.07 (-0.21 : 0.06) | -0.05 (-0.31 : 0.22) | -0.08 (-0.67 : 0.5) | 0.15 (0.04 : 0.27) |
| *Anthoxanthum odoratum* | 0.3 (-0.58 : 1.19) | 0.01 (-0.02 : 0.04) | -0.06 (-0.27 : 0.14) | -0.2 (-0.69 : 0.29) | 0.03 (-0.09 : 0.16) |
| *Anthyllis vulneraria* | 0.14 (-0.98 : 1.27) | -0.16 (-0.24 : -0.08) | -0.01 (-0.27 : 0.24) | 0.37 (-0.09 : 0.84) | 0.06 (-0.06 : 0.19) |
| *Brachypodium pinnatum* | -0.14 (-0.46 : 0.19) | 0.14 (0.12 : 0.16) | 0.02 (-0.16 : 0.2) | -0.01 (-0.44 : 0.41) | -0.33 (-0.43 : -0.23) |
| *Bromus hordeaceus* | 0.2 (-1.01 : 1.42) | -0.01 (-0.14 : 0.11) | -0.02 (-0.28 : 0.25) | -0.06 (-0.67 : 0.55) | 0.02 (-0.1 : 0.15) |
| *Bromus japonicus* | -0.06 (-0.38 : 0.25) | 0.12 (0.09 : 0.15) | 0.1 (-0.07 : 0.27) | -0.09 (-0.4 : 0.21) | -0.17 (-0.24 : -0.1) |
| *Centaurea nervosa* | 0.06 (-0.87 : 1) | -0.02 (-0.07 : 0.03) | 0.05 (-0.16 : 0.25) | 0.1 (-0.4 : 0.59) | 0.15 (0.01 : 0.29) |
| *Cynosurus cristatus* | -0.18 (-0.53 : 0.17) | 0.1 (0.08 : 0.13) | -0.02 (-0.2 : 0.16) | 0 (-0.46 : 0.45) | 0.02 (-0.05 : 0.09) |
| *Cynosurus echinatus* | 0.38 (-0.54 : 1.3) | 0.04 (-0.05 : 0.13) | 0 (-0.23 : 0.23) | -0.12 (-0.68 : 0.43) | 0.03 (-0.07 : 0.12) |
| *Dactylis glomerata* | 0.16 (-0.32 : 0.63) | 0.11 (0.08 : 0.14) | -0.06 (-0.25 : 0.14) | -0.16 (-0.59 : 0.26) | 0 (-0.1 : 0.09) |
| *Dianthus stenopetalus* | -0.26 (-1.42 : 0.91) | 0.09 (-0.01 : 0.19) | -0.05 (-0.3 : 0.21) | -0.08 (-0.64 : 0.48) | -0.1 (-0.21 : 0.01) |
| *Digitalis laevigata* | -0.29 (-1.46 : 0.88) | 0.08 (-0.02 : 0.19) | 0.18 (-0.07 : 0.43) | 0.2 (-0.36 : 0.77) | -0.03 (-0.16 : 0.11) |
| *Dorycnium germanicum* | -0.26 (-0.9 : 0.37) | -0.04 (-0.09 : 0) | -0.05 (-0.25 : 0.14) | 0.03 (-0.42 : 0.49) | -0.04 (-0.22 : 0.15) |
| *Festuca marginata subsp Heldreichii* | -0.01 (-0.39 : 0.37) | 0.02 (0 : 0.05) | -0.13 (-0.31 : 0.05) | -0.32 (-0.56 : -0.07) | -0.21 (-0.26 : -0.16) |
| *Festuca polita* | 0.9 (0.11 : 1.69) | 0.14 (0.1 : 0.18) | -0.09 (-0.29 : 0.12) | -0.38 (-0.84 : 0.09) | 0.01 (-0.08 : 0.09) |
| *Hordeum bulbosum* | 0.31 (-0.54 : 1.16) | 0.25 (0.21 : 0.28) | 0.22 (0.01 : 0.42) | -0.12 (-0.58 : 0.35) | -0.02 (-0.11 : 0.08) |
| *Leontodon crispus* | -0.19 (-1.23 : 0.85) | 0.08 (-0.05 : 0.21) | 0.03 (-0.24 : 0.29) | 0.33 (0.02 : 0.65) | 0.07 (-0.12 : 0.26) |
| *Lolium perenne* | -0.55 (-1.09 : -0.02) | 0.09 (0 : 0.17) | 0.05 (-0.21 : 0.31) | -0.08 (-0.49 : 0.32) | -0.01 (-0.1 : 0.08) |
| *Lotus corniculatus* | -0.22 (-0.88 : 0.44) | -0.14 (-0.17 : -0.11) | -0.06 (-0.26 : 0.13) | -0.03 (-0.34 : 0.27) | 0.14 (0.08 : 0.2) |
| *Melilotus indicus* | -0.02 (-1.21 : 1.17) | -0.05 (-0.17 : 0.08) | 0 (-0.26 : 0.26) | 0 (-0.6 : 0.61) | 0 (-0.13 : 0.12) |
| *Phleum pratense* | 0.21 (-0.33 : 0.75) | -0.03 (-0.11 : 0.04) | -0.09 (-0.29 : 0.11) | 0.53 (0.12 : 0.93) | -0.06 (-0.16 : 0.03) |
| *Pilosella leucopsilon* | 0.26 (-0.38 : 0.89) | -0.2 (-0.28 : -0.12) | -0.04 (-0.3 : 0.21) | 0.08 (-0.21 : 0.37) | 0.07 (-0.02 : 0.15) |
| *Piptatherum holciforme* | 0.04 (-0.35 : 0.43) | 0.36 (0.31 : 0.41) | 0.16 (-0.03 : 0.36) | -0.09 (-0.58 : 0.41) | -0.11 (-0.23 : 0.01) |
| *Plantago argentea* | -0.54 (-1.51 : 0.42) | 0 (-0.04 : 0.05) | 0.13 (-0.08 : 0.34) | 0.41 (-0.09 : 0.91) | 0.04 (-0.12 : 0.2) |
| *Plantago holosteum* | -0.16 (-1.41 : 1.09) | -0.01 (-0.13 : 0.11) | -0.06 (-0.32 : 0.2) | 0.25 (-0.09 : 0.6) | -0.01 (-0.2 : 0.17) |
| *Plantago lanceolata* | -0.01 (-0.92 : 0.9) | 0.02 (-0.04 : 0.07) | 0.11 (-0.1 : 0.33) | 0.29 (-0.14 : 0.72) | 0.05 (-0.08 : 0.18) |
| Species | SLA | Height | LA | LT | LDMC |
| *Potentilla recta* | 0.12 (-1.11 : 1.35) | 0.03 (-0.08 : 0.14) | 0.08 (-0.17 : 0.33) | 0.02 (-0.39 : 0.42) | -0.01 (-0.11 : 0.1) |
| *Rostraria cristata* | -1.18 (-1.89 : -0.46) | -0.08 (-0.17 : 0) | -0.09 (-0.31 : 0.12) | -0.27 (-0.72 : 0.18) | -0.06 (-0.15 : 0.03) |
| *Stachys germanica* | 0.18 (-0.75 : 1.1) | 0.04 (-0.07 : 0.16) | 0.13 (-0.11 : 0.37) | 0.02 (-0.57 : 0.6) | 0 (-0.15 : 0.14) |
| *Teucrium chamaedrys* | -0.18 (-1.4 : 1.05) | -0.1 (-0.25 : 0.04) | -0.03 (-0.3 : 0.24) | 0.07 (-0.33 : 0.48) | -0.12 (-0.28 : 0.04) |
| *Thymus longicaulis* | 1.72 (0.73 : 2.7) | -0.15 (-0.22 : -0.09) | -0.09 (-0.31 : 0.14) | -0.05 (-0.4 : 0.29) | 0.12 (0.03 : 0.21) |
| *Trifolium campestre* | 0.03 (-1.2 : 1.25) | -0.09 (-0.15 : -0.03) | -0.12 (-0.35 : 0.11) | -0.2 (-0.72 : 0.33) | 0.02 (-0.13 : 0.16) |
| *Trifolium fragiferum* | 0.28 (-0.89 : 1.44) | -0.11 (-0.23 : 0) | -0.03 (-0.28 : 0.23) | -0.08 (-0.67 : 0.5) | 0.07 (-0.06 : 0.2) |
| *Trifolium nigrescens* | -0.34 (-1.12 : 0.44) | -0.14 (-0.18 : -0.1) | -0.07 (-0.24 : 0.1) | 0.04 (-0.25 : 0.34) | 0.07 (-0.02 : 0.16) |
| *Trifolium parnassii* | -0.48 (-1.08 : 0.13) | -0.08 (-0.1 : -0.05) | 0.06 (-0.14 : 0.26) | -0.16 (-0.59 : 0.27) | 0.05 (-0.03 : 0.12) |
| *Viola macedonica* | -0.01 (-1.28 : 1.25) | -0.06 (-0.22 : 0.09) | 0 (-0.28 : 0.27) | -0.01 (-0.59 : 0.58) | 0.07 (-0.05 : 0.18) |

*Table S1 Species-specific interaction effects between aridity and grazing intensity for five functional traits (SLA, Height, LA, LT, LDMC). The values represent the estimated random slopes (coefficients) from the mixed-effects models, with 95% confidence intervals provided in parentheses. Shaded cells denote interactions where the confidence interval does not overlap with zero, indicating a statistically significant species-specific response to the combined pressure of aridity and grazing. Only species with at least three individual measurements per trait are included to ensure robust estimates.*
